# ProteinMCP: An Agentic AI Framework for Autonomous Protein Engineering

**DOI:** 10.64898/2026.03.11.711149

**Authors:** Xiaopeng Xu, Chenjie Feng, Chao Zha, Wenjia He, Maolin He, Bin Xiao, Xin Gao

## Abstract

Computational protein design is often constrained by slow, complex, inaccessible, and highly sophiscated and expert-dependent workflows that hinder its transferrability and generalization power for broader applications. We present ProteinMCP, an agentic AI framework designed to accelerate and democratize protein engineering. ProteinMCP automates end-to-end scientific tasks, delivering dramatic gains in efficiency; for instance, a comprehensive protein fitness modeling workflow was completed in just 11 minutes. This performance is achieved by an AI agent that intelligently orchestrates a unified ecosystem of 38 specialized tools, made accessible through a Model-Context-Protocol (MCP). A cornerstone of the framework is an automated pipeline that converts existing software into MCP-compliant servers, ensuring the platform is both powerful and perpetually extensible. We further demonstrate its capabilities through the successful autonomous design and selection of high-affinity de novo binders and therapeutic nanobodies. By removing technical barriers, ProteinMCP has the potential to shorten the design-build-test cycle and make advanced computational protein design accessible to the broader scientific community.

## Introduction

Computational protein design has emerged as a transformative discipline, enabling the creation of novel proteins with tailored functions and properties^1^. The ability to design proteins from first principles has led to breakthroughs in areas such as therapeutics, diagnostics, and biocatalysis^2–5^. Pioneers in this field have developed a suite of powerful computational tools, including the Rosetta software and more recent deep learning methods like RFdiffusion, that have made many of these advances possible^2,6,7^.

However, the rapid proliferation of these specialized tools has paradoxically created a major bottleneck. A typical design project now involves a complex, multi-step workflow that requires orchestrating numerous disparate software packages—each with its own unique interface and data formats—for tasks like multiple sequence alignment (MSA), structure prediction, and fitness modeling. This fragmentation of the toolchain is not only time-consuming and error-prone, but also demands a high level of computational expertise, effectively placing many advanced methods beyond the reach of the broader scientific community.

To address this challenge, we developed ProteinMCP, an agentic AI framework for autonomous protein engineering. ProteinMCP reimagines the protein design process by leveraging a Large Language Model (LLM) to interpret high-level scientific goals and autonomously execute complex workflows. The system integrates a comprehensive collection of 38 state-of-the-art protein design tools into a single, unified ecosystem through the Model-Context-Protocol (MCP), a standardization that enables seamless communication between the AI agent and the specialized software. A key innovation of our platform is an automated workflow that can create and deploy new MCP servers directly from existing code repositories, ensuring that ProteinMCP can be rapidly and sustainably expanded as new methods emerge.

In this manuscript, we present the architecture of ProteinMCP and demonstrate its capabilities through three case studies that represent common, yet complex, protein engineering challenges: protein fitness prediction, de novo binder design, and nanobody engineering. We show that ProteinMCP can automate these entire workflows, from initial data input to final analysis, with minimal human intervention. By drastically reducing execution times—for instance, completing a comprehensive fitness modeling analysis in just 11 minutes—and by systematically navigating complex design spaces, ProteinMCP empowers researchers to accelerate the design-build-test cycle. Ultimately, this work aims to democratize computational protein design, making powerful tools accessible to a wider range of researchers and enabling a new era of high-throughput, AI-driven protein engineering.

## Results

### The ProteinMCP Architecture: An Ecosystem for Agentic Protein Engineering

The architecture of ProteinMCP is a modular, four-tiered system designed to enable autonomous, AI-driven research (**Figure *1***). At its core, the Orchestration Layer contains a Large Language Model (LLM) agent that interprets high-level user goals, formulates plans, and executes them by invoking tools from the underlying MCP Server Layer. This server layer unifies a diverse and extensible ecosystem of 38 specialized bioinformatics tools—from sequence analysis to protein design—through a standardized Model-Context-Protocol (MCP). This design allows the agent to seamlessly integrate disparate software into complex workflows, transforming a collection of standalone programs into a cohesive and intelligent research platform.

**Figure 1.**
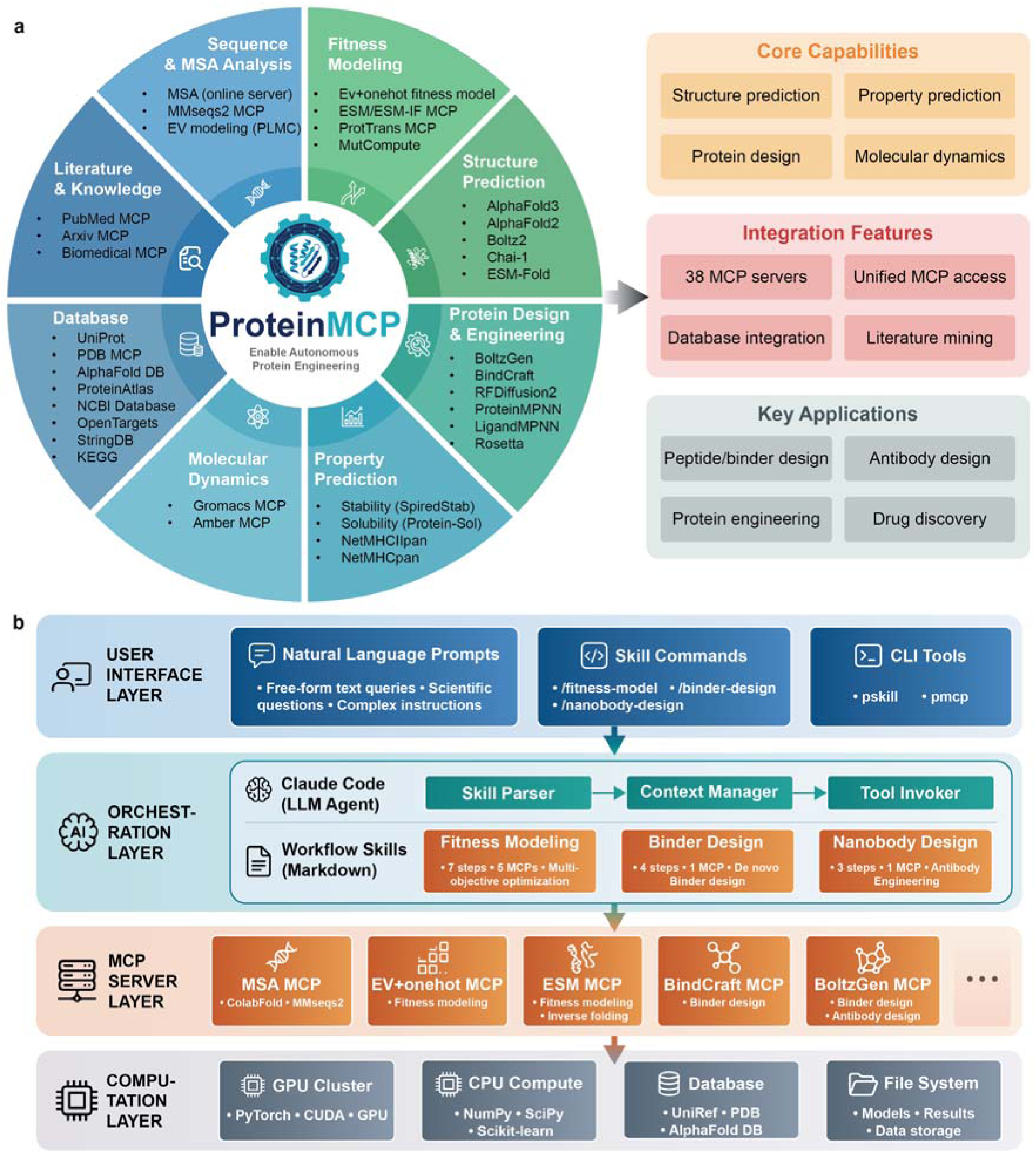
Overview of the ProteinMCP System Architecture and Capabilities. (a) The ProteinMCP ecosystem for autonomous protein engineering. The central platform integrates eight key modules: literature and knowledge query, sequence and multiple sequence alignment (MSA) analysis, fitness modeling, structure prediction, protein design and engineering, property prediction, molecular dynamics, and databases. Each module is equipped with a suite of specialized tools and Model-Context-Protocol (MCP) servers. The platform’s core capabilities include structure prediction, property prediction, protein design, and molecular dynamics. It features the integration of 38 MCP servers, unified MCP access, extensive database integration, and literature mining. Key applications of ProteinMCP encompass peptide and binder design, antibody design, general protein engineering, and drug discovery. (b) The multi-layered architecture of ProteinMCP. The system is composed of four distinct layers. The User Interface Layer allows users to interact with the system via natural language prompts, skill commands, or command-line interface (CLI) tools. The Orchestration Layer features an LLM agent (Claude Code) that interprets user requests, parses them into executable workflow skills, and manages the overall task execution through a Skill Parser, Context Manager, and Tool Invoker. The MCP Server Layer consists of numerous specialized servers that perform specific computational tasks, such as MSA, fitness modeling, and binder design. Finally, the Computation Layer provides the foundational resources, including GPU and CPU clusters, databases like UniRef and PDB, and a file system for storing models, results, and data.

### Automated Creation and Management of MCP Servers

A key bottleneck in building agentic systems for scientific research is the technical barrier to integrating existing tools. To overcome this, we developed an automated workflow for creating and deploying new MCP servers from existing code repositories (**Figure *2***). This process requires only the repository URL and the target functions as input. The system automatically sets up an isolated Conda environment, clones the repository, wraps the specified functions using our FastMCP library, and registers the new server with the host agent (**Figure S1**). This automated procedure drastically reduces the engineering effort required to expand the system’s capabilities. We have demonstrated its robustness by successfully converting 38 widely used bioinformatics tools into stable MCP servers, with nearly 100% success rate in MCP convertion, and very high success rate in executing complex, multi-step workflows. This streamlined process ensures that ProteinMCP can remain at the cutting edge by rapidly incorporating new tools and methods as they are developed by the scientific community.

**Figure 2.**
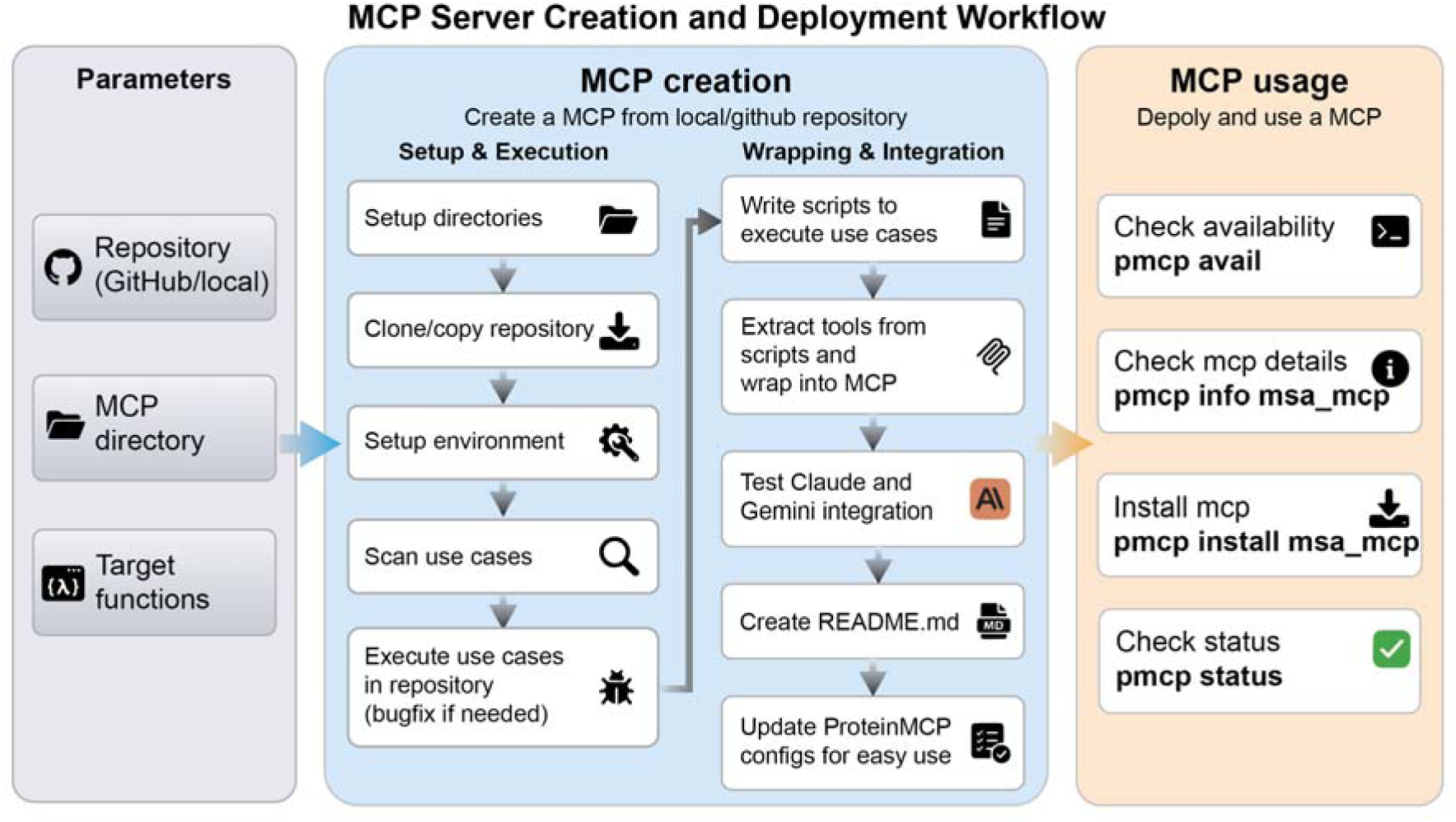
The Automated MCP Server Creation and Deployment Workflow in ProteinMCP. The workflow for automatically creating and deploying a new MCP server from an existing local or GitHub repository is a three-stage process. **Parameter Specification:** The process begins with the user providing three key parameters: the source code Repository (either a local path or a GitHub URL), the target MCP directory where the new server will be created, and the specific Target functions within the repository that are to be wrapped into MCP tools. **MCP Creation:** This central stage is divided into two parallel sub-workflows: 1) Setup & Execution: This sub-workflow prepares the environment by setting up the necessary directories, cloning or copying the repository content, establishing the required computational environment, scanning for use cases, and executing them to ensure functionality, with bug fixes applied as needed. 2) Wrapping & Integration: Concurrently, this sub-workflow focuses on integrating the code into the ProteinMCP ecosystem. It involves writing execution scripts for the identified use cases, extracting the core logic into MCP tools, testing the integration with LLM agents, automatically generating a README.md file for documentation, and finally, updating the ProteinMCP configuration files to make the new MCP easily accessible. **MCP Usage:** Once the MCP is created and integrated, it can be deployed and managed using the pmcp command-line tool. This includes checking for its availability (pmcp avail), inspecting its details (pmcp info <MCP_NAME>), installing it into the local environment (pmcp install <MCP_NAME>), and verifying its operational status (pmcp status).

### Comparative Analysis of Agentic Platforms

To position ProteinMCP in the context of the rapidly evolving landscape of AI agents for science, we conducted a qualitative comparison with four other recently published platforms: BioinfoMCP^8^, Paper2Agent^9^, PRIME^10^, and Biomni^11^ (**Table *1***). ProteinMCP is the only platform to achieve the highest rating (★★★) across all evaluated criteria, including ease of use, time savings, robustness, MCP creation, skill support, and token efficiency. While other platforms excel in specific areas, such as the automated conversion of papers to MCPs (Paper2Agent) or general-purpose biomedical task execution (Biomni), ProteinMCP distinguishes itself through its unique combination of a fully-featured, open-source framework, robust support for automated MCP creation, efficient token utilitization by supporting skill, and demonstrated high performance on complex, real-world protein engineering workflows. This positions ProteinMCP as a uniquely powerful and accessible platform designed by and for the protein science community to accelerate molecular engineering.

**Table 1.** Qualitative comparison of ProteinMCP against other state-of-the-art AI agent platforms in bioinformatics.

| Methods | Easy to master | Save time | Robustness | MCP creation | Workflow support | Token efficiency |
| --- | --- | --- | --- | --- | --- | --- |
| ProteinMCP | ??? | ??? | ??? | ??? | ??? | ??? |
| BioinfoMCP | ??? | ??? | ??? | ??? | ? | ?? |
| Paper2Agent | ?? | ??? | ??? | ??? | ? | ?? |
| PRIME | ?? | ??? | ??? | ? | ?? | ?? |
| Biomni | ??? | ??? | ??? | ? | ?? | ?? |

### Case Study 1: A Fully Automated and High-Throughput Fitness Modeling Workflow

Predicting the fitness of protein variants is a cornerstone of modern protein engineering. To demonstrate the power of ProteinMCP in this domain, we executed a fully automated workflow to build and evaluate a suite of fitness models for a target protein (**Figure *3*a**). Given only a wild-type sequence and a CSV file of variants with experimental fitness scores, the ProteinMCP agent autonomously executed a six-step workflow. This workflow encompassed both co-evolutionary models (MSA, PLMC^12^, EV+OneHot^13^) and models based on pre-trained protein language models (ESM^14,15^ and ProtTrans^16^), followed by automated aggregation and visualization of the results (**Figure S2**).

**Figure 3.**
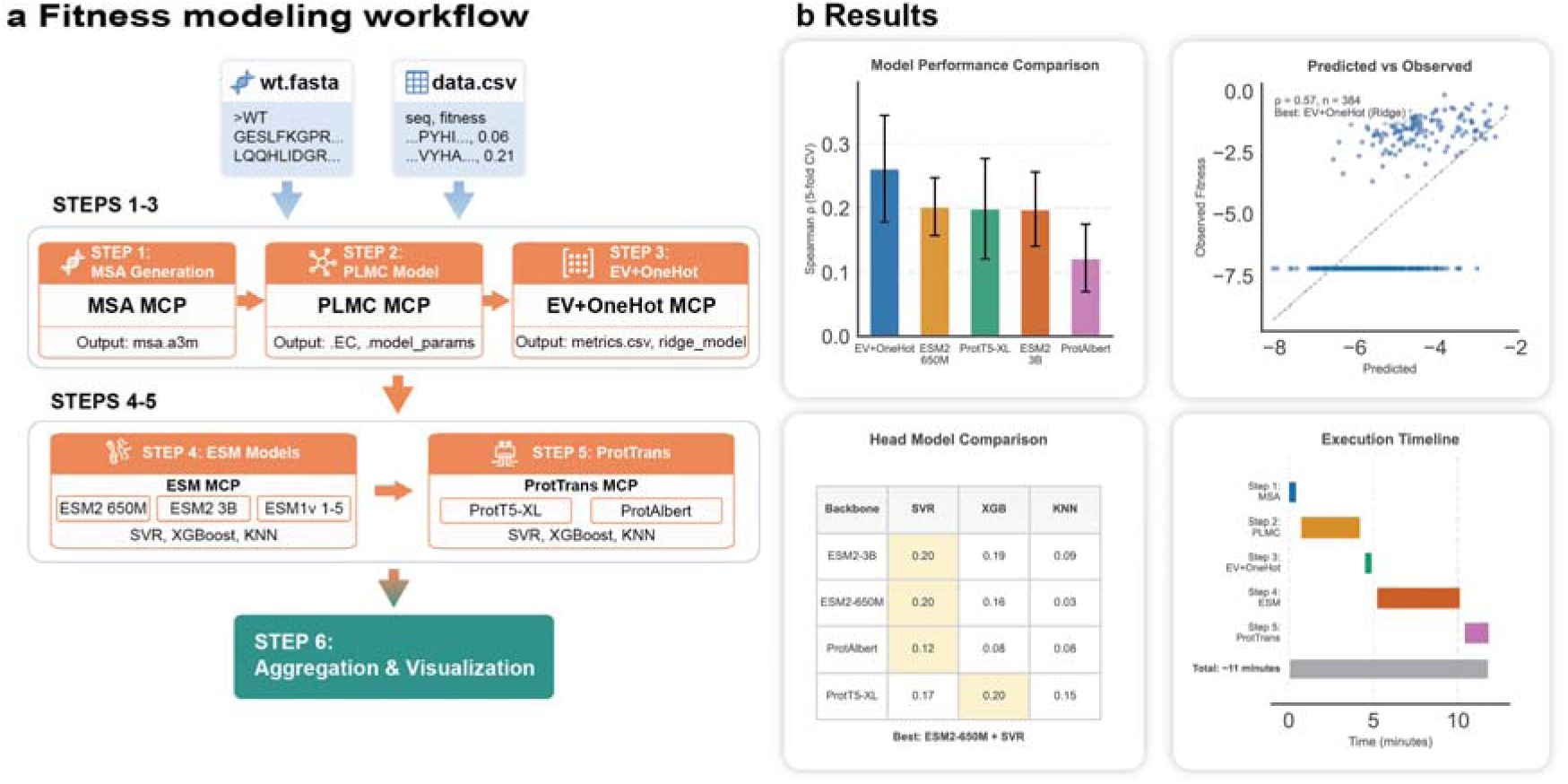
The Automated Fitness Modeling Workflow and Performance Analysis in ProteinMCP. (a) Fitness Modeling Workflow. The workflow is a six-step process that integrates multiple MCP servers to train and evaluate various fitness prediction models. Inputs: The workflow starts with two input files: wt.fasta, containing the wild-type protein sequence, and data.csv, which provides a list of sequence variants and their corresponding experimental fitness values. Steps 1-3: The initial steps focus on co-evolutionary models. Step 1 involves MSA generation using the MSA MCP. Step 2 utilizes the PLMC MCP to build a Potts model. Step 3 employs the EV+OneHot MCP to create a supervised fitness model based on evolutionary couplings and one-hot encoding. Steps 4-5: These steps leverage pre-trained protein language models. Step 4 uses the ESM MCP to evaluate various ESM) models, while Step 5 uses the ProtTrans MCP to evaluate models from the ProtTrans family (e.g., ProtT5-XL, ProtAlbert). Both steps pair the language model embeddings with supervised machine learning algorithms like SVR, XGBoost, and KNN. Step 6: The final step, Aggregation & Visualization, collects the outputs from all preceding models, compares their performance, and generates a consolidated results summary. (b) Results. This panel displays the aggregated and visualized outcomes of the workflow execution. Model Performance Comparison: A bar chart compares the predictive performance (Spearman’s ρ from 5-fold cross-validation) of the top models. In this example, the EV+OneHot model demonstrates the highest predictive accuracy. Predicted vs Observed Plot: A scatter plot shows the correlation between the predicted and observed fitness values for the best-performing model, EV+OneHot (Ridge), achieving a Spearman’s ρ of 0.57. Head Model Comparison: A table provides a detailed performance breakdown for different combinations of language model backbones and supervised learning heads, identifying ESM2-650M + SVR as another strong performer. Execution Timeline: A Gantt chart visualizes the execution time for each step, indicating that the entire workflow, encompassing the evaluation of multiple complex models, completes in approximately 11 minutes.

The entire process, which involved running multiple distinct and computationally intensive modeling pipelines, was completed in just 11 minutes (**Figure *3*b**). The system automatically identified the EV+OneHot model as the top performer, achieving a Spearman’s ρ of 0.57. The scientific impact of this automation is profound; it enables researchers to move from raw experimental data to a robust, cross-validated, and comparative assessment of multiple state-of-the-art fitness models in minutes, rather than days or weeks. This rapid, hands-off approach allows for high-throughput screening of modeling strategies, ensuring that researchers can quickly identify the optimal predictive model for their specific engineering objective, thereby accelerating cycles of design and testing.

To place this performance in context, accomplishing the same workflow manually would require an estimated one to three days of dedicated effort from a skilled bioinformatician. This conventional approach involves a series of time-consuming and error-prone tasks, including the individual setup and configuration of each software package, the development and debugging of custom integration scripts, the sequential execution and monitoring of each step, and the final aggregation and analysis of disparate outputs. Therefore, the 11-minute execution by ProteinMCP represents not just an incremental speed-up but a fundamental shift in efficiency, reducing the barrier to entry and freeing researchers to focus on scientific questions rather than on complex software integration.

### Case Study 2: Automated De Novo Design and Analysis of High-Affinity Binders

Designing novel proteins that bind to a specific target is a central goal in therapeutics and diagnostics. We tasked ProteinMCP with the de novo design of a binder to a target protein, with PDL1 as the target. The workflow, orchestrated by the BindCraft ^3^ MCP, successfully generated a series of candidate binders (**Figure *4*a**).

**Figure 4.**
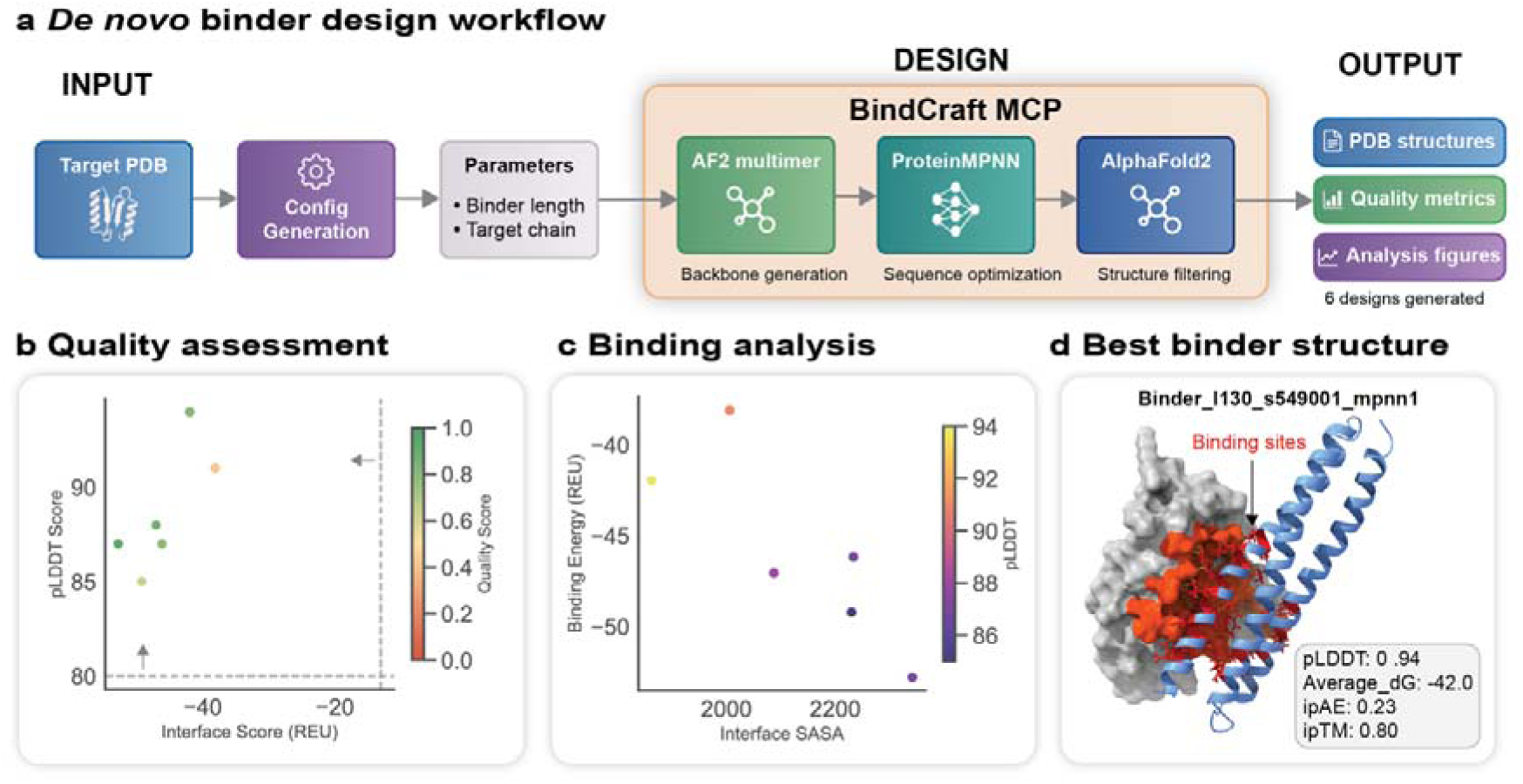
The Automated De Novo Binder Design and Analysis Workflow in ProteinMCP. (a) De Novo Binder Design Workflow. The process begins with user-defined inputs: a Target PDB structure and key Parameters such as the desired binder length and target chain, which are set during configuration generation. The core design process is orchestrated by the BindCraft MCP, which executes a three-step pipeline: 1) Backbone generation using AF2 multimer to create novel binder structures around the target, 2) Sequence optimization with ProteinMPNN to design optimal amino acid sequences for the generated backbones, and 3) Structure filtering with AlphaFold2 to predict the final complex structure and remove low-quality designs. The workflow’s Output consists of PDB structures for the designed binders, comprehensive quality metrics, and analysis figures, with six designs generated in this example. (b) Quality Assessment. This scatter plot provides an initial quality filter for the generated designs. Each point represents a single design, plotted by its Interface Score (REU, a measure of binding affinity) versus its pLDDT Score (a confidence metric for the predicted structure). Designs located in the upper-left quadrant (high pLDDT, low interface score) are considered high-quality candidates. The color of each point corresponds to a composite Quality Score. (c) Binding Analysis. Promising candidates are further evaluated in this plot, which shows the relationship between the predicted Binding Energy (REU) and the Interface Solvent Accessible Surface Area (SASA). Lower binding energy values indicate more favorable interactions. The points are color-coded by their pLDDT score, allowing for a multi-objective assessment of the designs’ stability and binding potential. (d) Best Binder Structure. The top-ranked design is visualized in complex with its target. The designed binder (blue helix) is shown interacting with the target protein (grey surface), with the specific binding sites highlighted in red. Key performance metrics for this design are displayed, including a pLDDT of 0.94, an average binding free energy (dG) of -42.0, an interface predicted Aligned Error (ipAE) of 0.23, and an interface predicted TM-score (ipTM) of 0.80, all of which indicate a high-confidence and high-affinity binder prediction.

Crucially, the binder design workflow did not stop at generation but proceeded to a multi-stage, automated analysis to identify the most promising candidates. After the initial “hallucination” of binder backbones and sequences using AlphaFold2 (AF2) multimer^17^, trajectories are rejected if they fail to meet baseline metrics for structural confidence (pLDDT > 0.7) or physical contacts. Successful trajectories then undergo MPNNsol^18^ sequence optimization to enhance stability and solubility while maintaining the designed interface.

These optimized designs are subjected to a rigorous in silico filtering process involving AF2 monomer^19^ reprediction to ensure a robust, unbiased assessment of the interface. The final quality assessment combines deep learning metrics—specifically pLDDT (> 0.8) and i_pTM (>0.5)—with Rosetta^6^ physics-based scoring, which evaluates interface shape complementarity and the number of unsaturated hydrogen bonds. The workflow typically culminates in the selection of the top-ranked binders (ranked by i_pTM) for experimental validation, which has yielded binders with nanomolar affinity across diverse and challenging targets.

The results of an automated design-and-filter process are visualized to provide a comprehensive overview of the candidate pool (**Figure 4b-d** and **Figure S3**). The quality assessment, based on a composite score of structural confidence (pLDDT) and binding affinity (Interface Score), allows for clear ranking of the designs (**Figure *4*b**, **Figure S3**). A more detailed binding analysis further characterizes the top candidates (**Figure *4*c**). This workflow culminated in the identification of a top-ranked binder with outstanding predicted metrics, including a pLDDT of 0.94 and an interface TM-score of 0.80, indicating a high-confidence prediction of a stable and specific interaction (**Figure *4*d**). By automating this intricate design-and-filter process, ProteinMCP provides a powerful, high-throughput platform for the rapid discovery of novel binders with therapeutic potential.

### Case Study 3: Autonomous Engineering and Selection of Nanobodies

Nanobodies represent a rapidly emerging class of therapeutics, but their engineering remains a complex, multi-parameter optimization problem. We deployed ProteinMCP to perform an end-to-end nanobody design workflow using the BoltzGen^20^ MCP. The system autonomously managed the entire process, from job configuration and submission to results retrieval and analysis.

A demo case is run with 50 designs (**Figure *5*a**). While the underlying BoltzGen tool employs a complex, native filtering pipeline to rank its 50 generated designs, ProteinMCP’s primary role is to make this powerful but specialized tool easily accessible and its outputs readily interpretable. The filtering within BoltzGen relies on a multi-metric evaluation, considering core metrics such as structural confidence (pTM), interface confidence (ipTM), predicted alignment error (pAE), and the number of hydrogen bonds at the interface.

**Figure 5.**
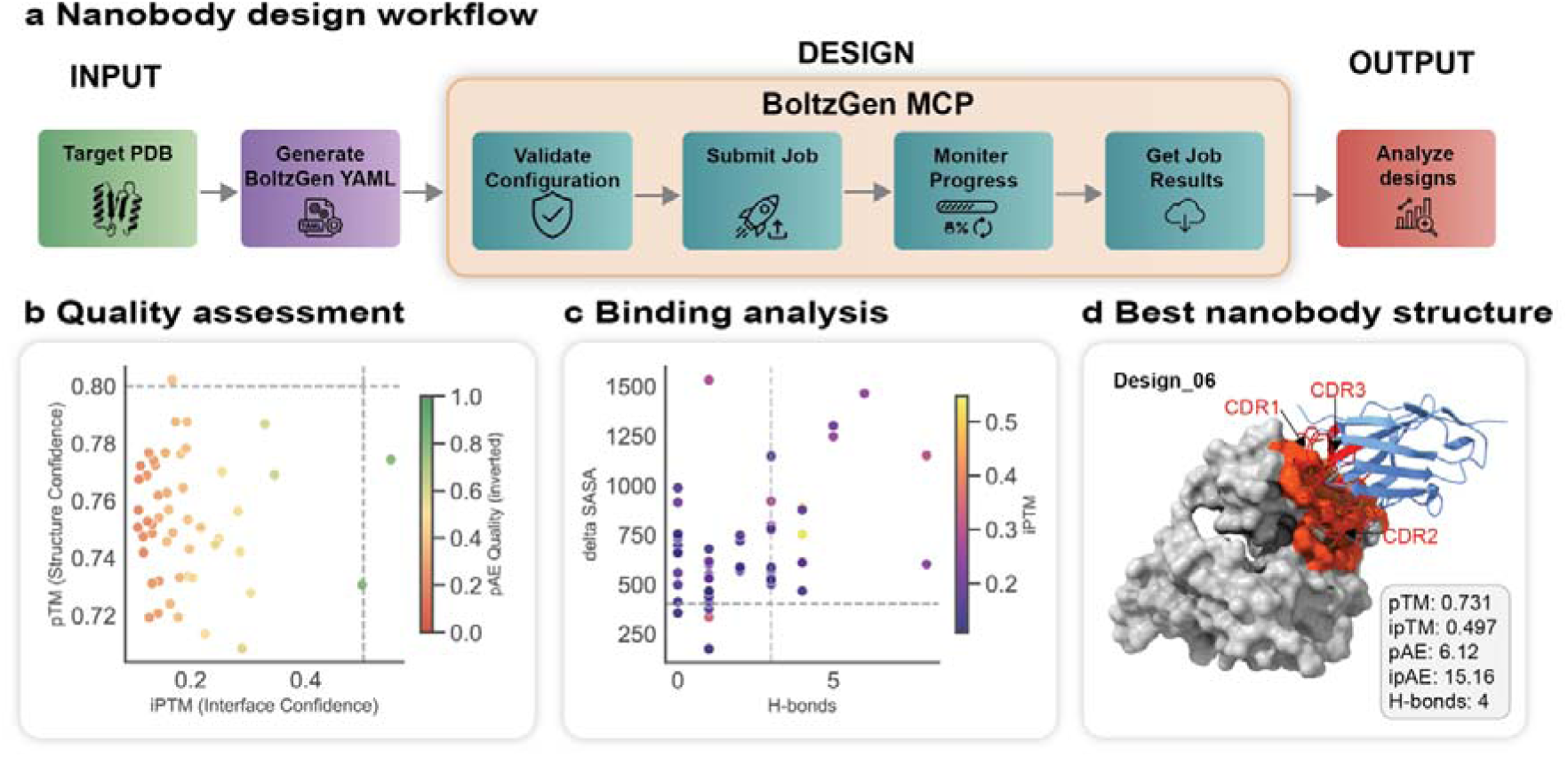
The Automated Nanobody Design and Analysis Workflow in ProteinMCP. (a) Nanobody Design Workflow. The process is initiated with a Target PDB structure. A BoltzGen yaml configuration file is then generated to specify the design parameters. The core design process is managed by the BoltzGen MCP, which follows a four-step procedure: Validate Configuration, Submit Job to the computational backend, Monitor Progress of the design calculations, and Get Job Results upon completion. The resulting designs are then passed to the Output stage for in-depth analysis. (b) Quality Assessment. This scatter plot serves as the primary filter for evaluating the generated nanobody designs. Each design is plotted based on its pTM (Structure Confidence) and ipTM (Interface Confidence) scores. Designs populating the upper-right quadrant, indicating high confidence in both the overall structure and the binding interface, are selected for further analysis. The color scale represents the pAE Quality (inverted), where greener points signify higher quality (lower predicted alignment error). (c) Binding Analysis. This plot provides a deeper analysis of the binding characteristics of the filtered candidates. It correlates the number of hydrogen bonds (H-bonds) formed at the interface with the change in solvent-accessible surface area (delta SASA) upon binding. A larger delta SASA and a higher number of H-bonds are indicative of a more extensive and specific binding interface. The points are color-coded by their ipTM score, reinforcing the focus on high-confidence interface predictions. (d) Best Nanobody Structure. The best-performing candidate, Design_06, is visualized in its bound state. The nanobody (blue cartoon) is shown interacting with the target protein (grey surface). The critical binding regions of the nanobody, the Complementarity-Determining Regions (CDR1, CDR2, CDR3), are highlighted in red, forming a precise interface with the target (orange). Key metrics quantifying the quality of this design are provided, including a pTM of 0.731, an ipTM of 0.497, and the formation of 4 hydrogen bonds, confirming a high-quality and specific interaction.

The comprehensive results from the automated workflow are summarized in a series of custom visualizations designed to facilitate rapid assessment of the candidate pool. The initial quality assessment plot (**Figure *5*b**) displays all 50 designs, mapping their overall structural confidence (pTM) against their interface confidence (ipTM). This visualization, colored by the predicted alignment error (pAE), clearly distinguishes the two high-quality designs that passed the stringent filtering from the 48 that failed. For a more granular view, the binding analysis plot (**Figure *5*c**) examines the relationship between the change in solvent-accessible surface area (delta SASA) and the number of hydrogen bonds, providing deeper insight into the physical interactions at the binding interface. Finally, the structure of the top-ranked candidate, Design_06, is presented in detail (**Figure *5*d**), highlighting its three complementarity-determining regions (CDRs) and a summary of its excellent predicted metrics (pTM: 0.731, ipTM: 0.497, H-bonds: 4). A more detailed breakdown of all metrics across the 50 designs is provided in **Figure S4**. This case study demonstrates ProteinMCP’s strength not in reinventing design algorithms, but in seamlessly orchestrating them and presenting their outputs in a clear, actionable format, thereby accelerating the discovery of high-potential therapeutic candidates.

## Discussion

The three case studies presented in this manuscript—fitness modeling, de novo binder design, and nanobody engineering—illustrate the transformative potential of ProteinMCP to accelerate and democratize computational protein engineering. By leveraging an agentic AI framework to orchestrate a comprehensive suite of state-of-the-art tools, ProteinMCP directly addresses the significant fragmentation that has traditionally hindered the field. The platform’s ability to autonomously interpret high-level goals and execute complex, multi-step workflows allows researchers to transcend the tedious and error-prone process of manual tool integration and focus instead on the core scientific questions that drive discoveries.

Our case studies highlight several key advantages of the ProteinMCP platform. First, the modularity of the MCP ecosystem enables the flexible combination of diverse tools to address a wide range of design challenges. The fitness modeling workflow, for example, seamlessly integrated and compared co-evolutionary methods with multiple protein language models, a task that would typically require extensive manual scripting and data wrangling. Unlike rigid, predefined pipelines where toolchains are hardcoded, the agentic orchestrator dynamically selected and sequenced these tools based on the evolving context of the analysis, showcasing a level of flexibility that is a significant conceptual advance. Second, the automation of the design-analyze-select cycle enables the rapid and rigorous exploration of a vast design space. This was particularly evident in the nanobody design case study, where the agent autonomously filtered an initial set of 50 candidates down to two high-quality designs based on a sophisticated, multi-metric analysis of structural integrity and binding interface properties. This autonomous, data-driven decision-making stands in stark contrast to conventional workflows, which would necessitate stopping the pipeline to manually inspect intermediate results and decide on the next steps. This ability to not only generate but also critically evaluate designs is a hallmark of a truly intelligent agent. Finally, the platform’s efficiency—demonstrated by the 11-minute execution of the entire fitness modeling pipeline—enables a pace of research that was previously unattainable. This dramatic speed-up is not merely a result of faster tools, but a direct consequence of the agentic model’s ability to eliminate the idle time and manual hand-offs that plague traditional, fragmented pipelines, thereby accelerating the entire design-build-test cycle.

The practical utility of a framework like ProteinMCP also depends on its robustness and ability to gracefully handle the inevitable errors that arise from a complex chain of integrated tools. The platform’s stability is achieved through several complementary mechanisms. At the infrastructure layer, each MCP server is encapsulated in an isolated runtime environment—either a local Python virtual environment or a Docker container—ensuring that failures in one tool do not propagate and destabilize the entire workflow. For computationally intensive tools, the use of stateless Docker containers that are created fresh for each invocation prevents state accumulation and resource leaks. At the orchestration layer, the framework delegates error handling to the AI agent, which can interpret error messages and autonomously attempt to resolve the issue, for instance by adjusting parameters or retrying a call. This agentic error recovery is a key advantage over traditional scripted pipelines, allowing the system to respond flexibly to the diverse failure modes of heterogeneous computational biology tools. Finally, the pmcp status command provides a pre-flight health-check mechanism, allowing users to verify that all registered MCPs are responsive before initiating a workflow, thereby reducing the likelihood of mid-workflow failures.

Furthermore, the automated MCP creation workflow is a critical innovation that ensures the long-term viability and relevance of the ProteinMCP ecosystem. Science evolves rapidly, and an agentic platform is only as powerful as the tools it can access. By providing a streamlined, automated process for integrating new tools, we have created a sustainable and extensible platform that can grow with the field. This stands in contrast to other platforms that may require significant manual effort to incorporate new functionalities, as shown in our comparative analysis (**Table *1***).

This inherent extensibility makes the framework readily applicable to other challenging domains, such as enzyme design and optimization. While a comprehensive enzyme design workflow is part of our future work, ProteinMCP is already well-prepared for this task. The platform’s modularity means that many foundational tools are already available as MCPs, and its architecture is explicitly designed to absorb new, specialized methods—such as Placer^21^ for placing catalytic motifs—as they emerge in this fast-evolving field. This ability to seamlessly integrate new capabilities ensures that ProteinMCP can adapt to orchestrate complex, evolving workflows, positioning it as a durable and versatile platform for future protein engineering challenges.

While ProteinMCP represents a significant step forward, it is important to acknowledge its limitations. The platform’s capabilities are inherently dependent on the quality and scope of the integrated tools. The success of any design project ultimately rests on the power of the underlying algorithms, and the agent cannot transcend their intrinsic limitations. Furthermore, all predictions made by ProteinMCP are computational and require experimental validation. Indeed, the dramatic increase in the throughput of computational design enabled by platforms like ProteinMCP underscores the urgent need for parallel advances in high-throughput experimental methods. Closing this “design-to-data” loop will be crucial for realizing the full potential of AI-driven protein engineering.

In conclusion, ProteinMCP provides a robust, extensible, and user-friendly platform that empowers researchers to tackle complex protein engineering challenges with unprecedented speed and autonomy. By shifting the paradigm from manual tool-chaining to high-level goal-oriented instruction, we believe ProteinMCP will not only accelerate discovery for experts but also make the power of computational protein design accessible to a broader scientific community, fostering a new era of innovation in molecular science.

## Methods

### Automated MCP Server Creation

The integration of new tools into the ProteinMCP ecosystem is facilitated by an automated MCP creation pipeline orchestrated by an LLM agent (Claude). This process, encapsulated within the MCPCreator class, transforms a standard software repository (either from a GitHub URL or a local path) into a fully compliant MCP server. The pipeline follows a structured, multi-step process (**Figure *2***): 1.Environment Setup: A dedicated directory for the new MCP server is created, including subdirectories for the cloned repository (repo/), the server source code (src/), and a sandboxed Conda virtual environment (env/). 2.Code Ingestion: The target software repository is either cloned from GitHub or copied from a local source into the repo/ directory. 3.Use-Case Identification: The LLM agent analyzes the repository’s documentation (e.g., README files, tutorials, and examples) to identify a set of primary use cases. This can be guided by a user-provided filter to focus on specific functionalities. 4.Functionality Verification: The agent attempts to execute these use cases directly, ensuring the underlying code is functional and identifying any immediate bugs or dependency issues. 5.Script Generation: For each verified use case, the agent writes a Python script that encapsulates the required function calls and logic. This step abstracts the core functionality into a set of executable scripts. 6.MCP Wrapping: The generated scripts are then wrapped into a standardized server interface using the FastMCP library. The agent defines tool endpoints corresponding to each use case, creating a server.py file that serves as the entry point for the MCP server. 7.Agent Integration Testing: The newly created MCP server is tested for integration with the host agent (Claude Code) to ensure seamless communication and execution. 8.Documentation Generation: Finally, the agent generates a comprehensive README.md file for the MCP server, documenting its capabilities, tool definitions, and usage examples.

This entire process is managed by the **pmcp create** command-line interface, which automates the LLM interactions and file system operations, enabling the rapid and scalable expansion of the ProteinMCP tool universe.

### Skill-Based Workflow Abstraction

While MCPs provide standardized interfaces to individual tools, complex scientific tasks require the orchestration of multiple tools in a specific sequence. In ProteinMCP, these multi-step procedures are abstracted into Skills. A Skill is a human-readable Markdown document that defines a complete scientific workflow, including its purpose, required MCPs, configuration parameters, and a sequence of execution steps.

The **pskill** command-line utility, powered by the SkillCreator and SkillManager classes, is used to manage these skills. The creation of a skill involves defining the workflow logic in a structured format. For example, the fitness_modeling.md skill specifies its reliance on the msa_mcp, plmc_mcp, esm_mcp, and prottrans_mcp servers and outlines the precise sequence of operations, from MSA generation to model training and final visualization.

This skill-based abstraction offers two key advantages. First, it makes complex workflows token-efficient. Instead of providing the LLM agent with lengthy, detailed instructions for every run, the user can simply invoke the skill (e.g., /fitness-model), and the agent executes the pre-defined, validated procedure. Second, it enhances reproducibility and reusability, as skills can be version-controlled, shared, and executed by different users with guaranteed consistency.

### Agentic Workflow Execution and Debugging

Workflows in ProteinMCP are executed by an LLM agent (Claude Code) that interprets the Skill Markdown files. The agent parses the document, understands the sequence of steps, and invokes the necessary MCP tools with the correct parameters, managing the data flow between steps.

Debugging in this agentic framework is an interactive and conversational process. If a workflow step fails or produces an unexpected result, the user can engage directly with the agent. The debugging process typically involves: 1.Observing the Failure: The agent provides detailed logs and error messages from the specific MCP that failed. 2.Instructing the Agent: The user can analyze the error and provide natural language instructions to the agent to fix it. For example, the user might instruct the agent to “modify the server.py file of the esm_mcp to handle empty sequences” or “adjust the parameters in Step 3 of the binder_design.md skill.” 3.Iterative Refinement: The agent applies the suggested changes to the underlying MCP server code or the Skill Markdown file and re-executes the failed step. This iterative, conversational loop continues until the bug is resolved and the workflow runs successfully.

This approach leverages the code generation and understanding capabilities of the LLM to create a powerful, interactive debugging environment that does not require the user to have deep expertise in the underlying codebase of each tool.

### Workflow Benchmarking and Case Study Implementation

The performance and capabilities of ProteinMCP were benchmarked through the implementation of three representative protein engineering workflows. The primary goal of this benchmarking was not to establish new state-of-the-art performance in the tasks themselves, but rather to qualitatively and persuasively demonstrate the platform’s ability to automate complex, end-to-end scientific tasks with significant gains in efficiency and accessibility.

For each case study, the workflow was executed from a high-level user prompt, and the agent autonomously performed all subsequent steps. Key performance indicators, such as total execution time, were recorded. The parameters for each workflow were specified in their respective skill files, including protein name, wt.fasta file path, data.csv file path and output directory for fitness modeling workflow; target pdb file, chain, binder length, number of designs, and output directory for binder design workflow; target cif file, chain, nanobody scaffolds, number of designs, and output directory for nanobody design workflow.

The results of these benchmarks are presented not just as raw data, but as comprehensive analysis figures automatically generated by the workflows themselves (**Figure *3*Figure *5***). This demonstrates the platform’s ability to not only execute the core computational tasks but also to perform the subsequent data analysis and visualization, delivering publication-ready insights directly to the user. This qualitative demonstration of end-to-end automation serves as a persuasive benchmark of the system’s scientific utility.

## Author Contributions

- Xiaopeng Xu: Conceptualization, Methodology, Software, Investigation, Data Curation, Writing – Original Draft, Visualization.
- Chenjie Feng: Writing – Review & Editing.
- Chao Zha: Investigation, Writing – Review & Editing.
- Wenjia He: Writing – Review & Editing.
- Maolin He: Writing – Review & Editing.
- Bin Xiao: Writing – Review & Editing.
- Xin Gao: Supervision, Project Administration, Funding Acquisition, Writing – Review & Editing.

## Supporting information

SI of ProteinMCP

## Acknowledgments

This publication is based upon work supported by Office of Research Administration (ORA) of the King Abdullah University of Science and Technology (KAUST) under Award No REI/1/5234-01-01, REI/1/5414-01-01, REI/1/5289-01-01, REI/1/5404-01-01, REI/1/5992-01-01, URF/1/4663-01-01, Center of Excellence for Smart Health (KCSH), under award number 5932, and Center of Excellence on Generative AI, under award number 5940, and the Natural Science Foundation of Ningxia Province [2024AAC03247, 2024BEH04022].

We acknowledge the computational resources of the Supercomputing Laboratory at King Abdullah University of Science & Technology (KAUST). We thank Bestzyme for providing the industrial context of this study. We thank the developers of the various open-source tools integrated into ProteinMCP for their contributions to the scientific community.

## Data Availability Statement

The ProteinMCP framework is publically available for academic use at https://github.com/charlesxu90/proteinmcp. All data and code used in the case studies are available within the platform and can be reproduced following the workflows described in this manuscript.

