## Supplementary material for "ProteinMCP: An Agentic AI Framework for Autonomous Protein Engineering": SI of ProteinMCP

Supplementary Information of ProteinMCP: An Agentic AI Framework for Autonomous Protein Engineering

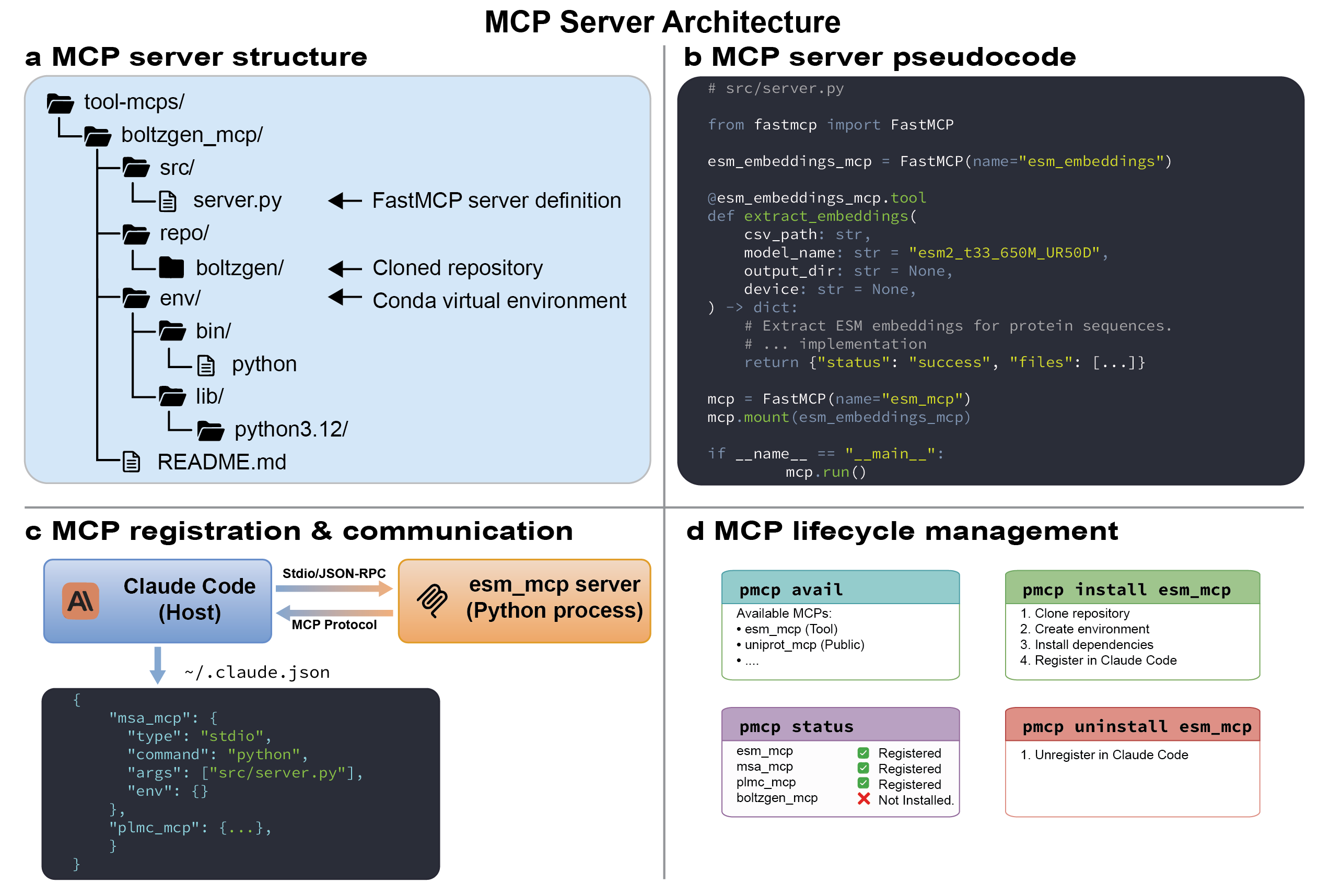

**Figure S1 | The MCP Server Architecture in ProteinMCP.** (a) MCP Server Structure. Each MCP server is encapsulated within a standardized directory structure. As shown for the boltzgen_mcp, this includes: the src directory containing the server.py definition; the repo directory holding a cloned version of the original tool's source code; the env directory containing a self-contained Conda virtual environment with all necessary dependencies; and a README.md file for documentation. This modular structure ensures that each tool is isolated and reproducible. (b) MCP Server Pseudocode. This panel illustrates the simplicity of converting a Python function into an MCP tool using the FastMCP library. A standard Python function, such as extract_embeddings, is decorated with @<mcp_instance_name>.tool. The function is then mounted onto a FastMCP instance, which handles the underlying server creation and communication. The mcp.run() command starts the server, making the tool accessible to the host agent. (c) MCP Registration & Communication. The host agent (Claude Code) communicates with MCP servers, which run as independent Python processes, using the MCP Protocol over a standard I/O (Stdio) JSON-RPC channel. MCPs are registered in a central ~/.claude.json configuration file. This file contains entries for each MCP, specifying the command and arguments required to launch it, thereby enabling the host to discover and invoke the tools as needed. (d) MCP Lifecycle Management. The pmcp command-line interface provides a suite of tools for managing the entire lifecycle of an MCP server. pmcp avail lists all available MCPs. pmcp install <mcp_name> automates the entire setup process, including cloning the repository, creating the environment, installing dependencies, and registering the MCP with the host. pmcp status checks the current registration and installation status of all MCPs, and pmcp uninstall <mcp_name> handles the removal and unregistration of an MCP.

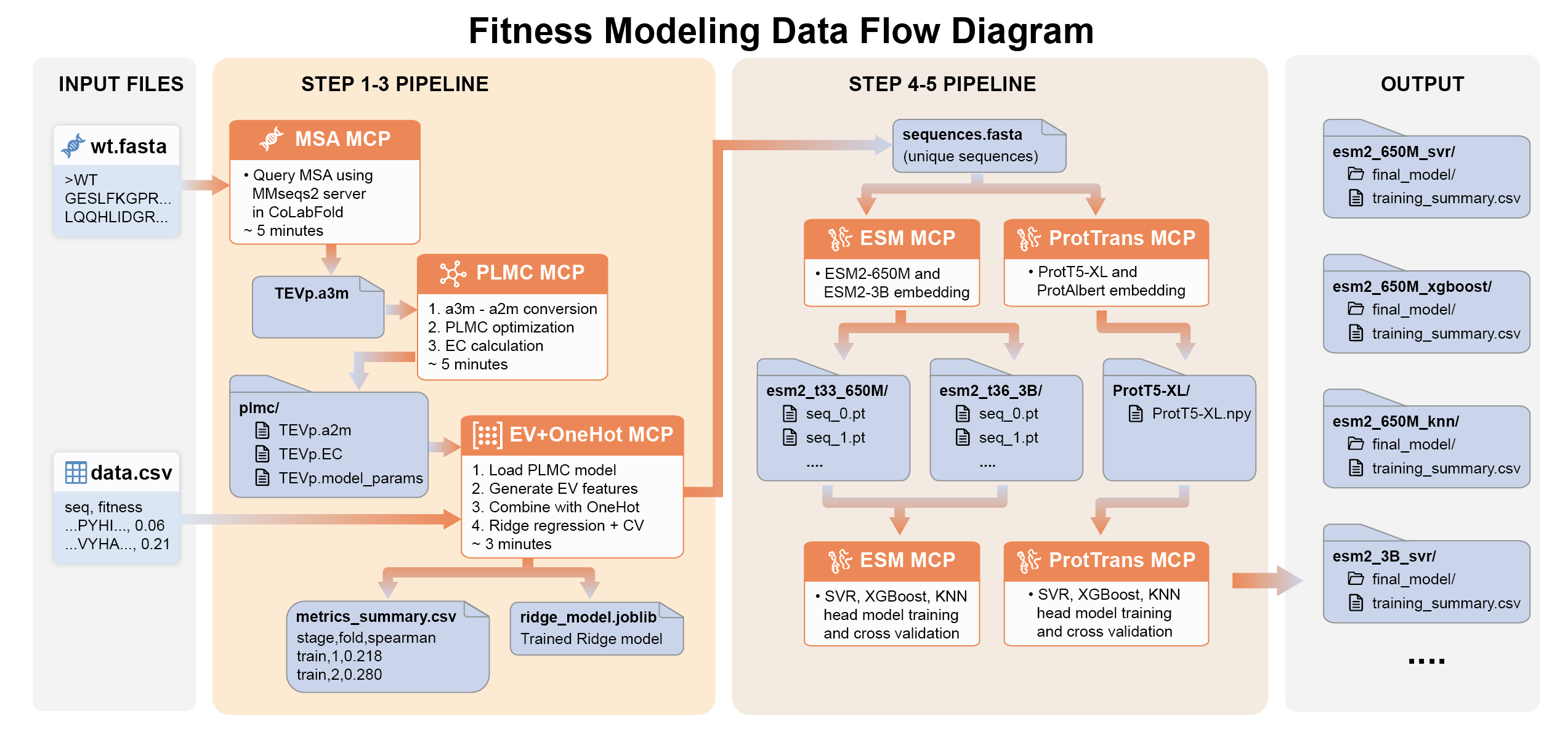

**Figure S2 | Detailed Data Flow of the Automated Fitness Modeling Workflow in ProteinMCP.** This diagram provides a granular view of the data flow and intermediate file generation within the fitness modeling workflow, illustrating how data is processed and passed between different MCPs. **Input Files:** The workflow is initiated with two files: wt.fasta, containing the wild-type protein sequence, and data.csv, a table of sequence variants and their experimentally measured fitness scores. **Step 1-3 Pipeline (Co-evolutionary Models):** This pipeline focuses on models derived from Multiple Sequence Alignments (MSAs). The wt.fasta file is fed into the MSA MCP, which uses the MMseqs2 server to generate an MSA file (TEVp.a3m); this MSA file is then processed by the PLMC MCP, which performs PLMC optimization and calculates evolutionary couplings (EC), outputting several files including the model parameters (TEVp.model_params) and ECs (TEVp.EC); the EV+OneHot MCP takes the PLMC model outputs along with the data.csv file to train a supervised model. It generates EV (evolutionary) features, combines them with a one-hot encoding of the sequences, and trains a Ridge regression model with cross-validation. This step produces a metrics_summary.csv file with performance data and a ridge_model.joblib file containing the trained model. **Step 4-5 Pipeline (Protein Language Models):** This pipeline runs in parallel and leverages large-scale, pre-trained protein language models. Unique sequences from data.csv are compiled into a sequences.fasta file. This file is passed to both the ESM MCP and the ProtTrans MCP. These MCPs generate embeddings for each sequence using various models (e.g., ESM2-650M, ESM2-3B, ProtT5-XL, ProtAlbert), resulting in sets of embedding files (e.g., .pt and .npy files). The generated embeddings are then used by the ESM MCP and ProtTrans MCP to train multiple supervised "head" models (SVR, XGBoost, KNN) for each embedding type, again using the fitness values from data.csv and performing cross-validation. **Output:** The workflow concludes by generating a structured set of output directories. For each combination of a language model (e.g., esm2_650M) and a head model (e.g., svr), a dedicated directory is created. Each directory contains the final_model/ and a training_summary.csv file, providing a comprehensive and organized summary of all trained models and their performance.

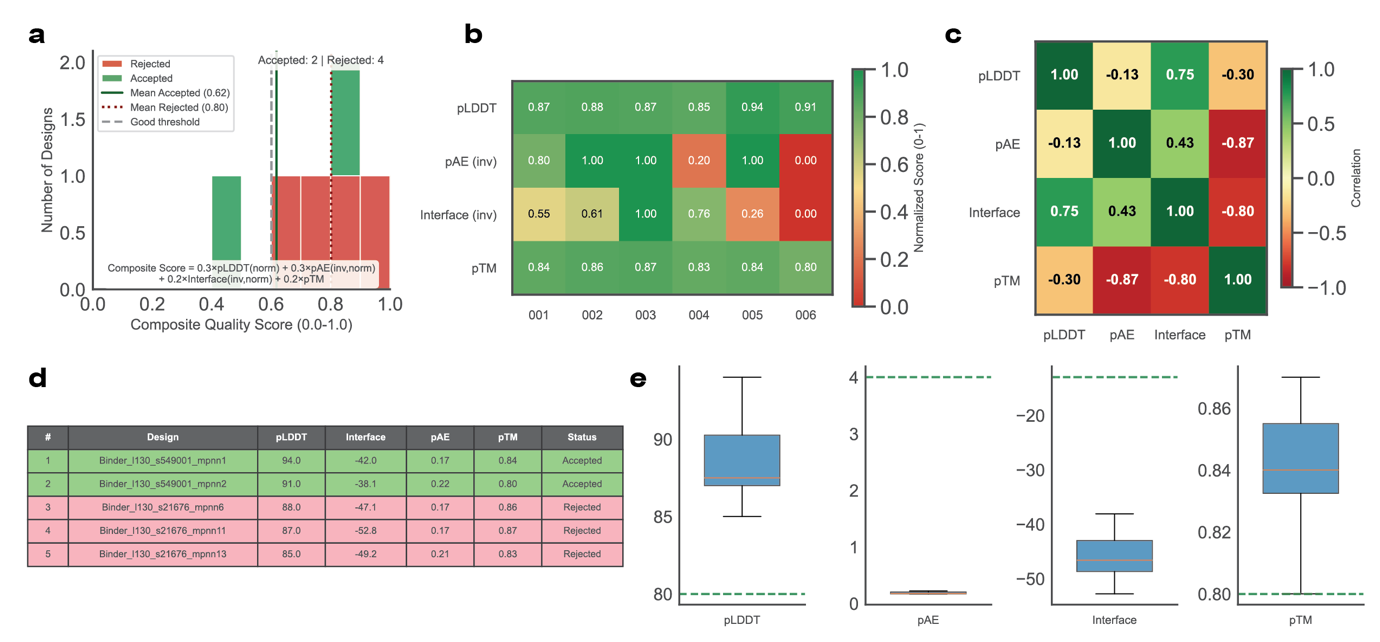

**Figure S3 | Detailed Analysis of Binder Design Metrics and Selection Criteria in ProteinMCP.** This figure presents a multi-faceted analysis of the metrics used to evaluate and select the de novo designed binders, providing a deeper insight into the quality control process. (a) Composite Quality Score Distribution. This histogram displays the distribution of a composite quality score, which consolidates multiple performance indicators into a single value. The score is a weighted sum of normalized pLDDT, pAE (inverted), Interface score (inverted), and pTM. The plot clearly distinguishes between 'Accepted' (green) and 'Rejected' (red) designs, showing a significant separation between the mean scores of the two groups (0.80 for accepted vs. 0.62 for rejected) relative to a predefined quality threshold. (b) Per-Design Normalized Score Matrix. This heatmap provides a normalized (0-1) overview of each design's performance across four key metrics: pLDDT (structure confidence), pAE (interface alignment error, inverted), Interface score (binding affinity, inverted), and pTM (interface confidence). The color gradient from red (poor) to green (excellent) allows for a quick comparative assessment, highlighting that the accepted designs (e.g., 001, 002) consistently score well across most metrics, whereas rejected designs (e.g., 005, 006) fail on one or more key indicators. (c) Inter-Metric Correlation Analysis. This correlation matrix reveals the relationships between the different quality metrics. Strong positive correlations are observed between pLDDT and the Interface score (0.75), indicating that well-structured designs tend to have better binding interfaces. Conversely, a strong negative correlation is seen between pAE and pTM (-0.87), which is expected as lower alignment errors at the interface correspond to higher interface confidence scores. (d) Top-Ranked Design Candidates. This table presents a quantitative summary of the top five designs, listing their raw scores for pLDDT, Interface, pAE, and pTM, along with their final 'Accepted' or 'Rejected' status. This provides the specific data points that underpin the visual analyses in the other panels. (e) Statistical Distribution of Design Metrics. This series of box plots visualizes the statistical distribution (median, quartiles, and range) of the raw scores for each of the four key metrics across all generated designs. The dashed green lines indicate the thresholds for good performance, providing context for the overall quality of the design batch. For instance, the majority of designs surpassed the pLDDT threshold of 80 and the pAE threshold of 4.0, but showed wider variance in Interface scores and pTM.

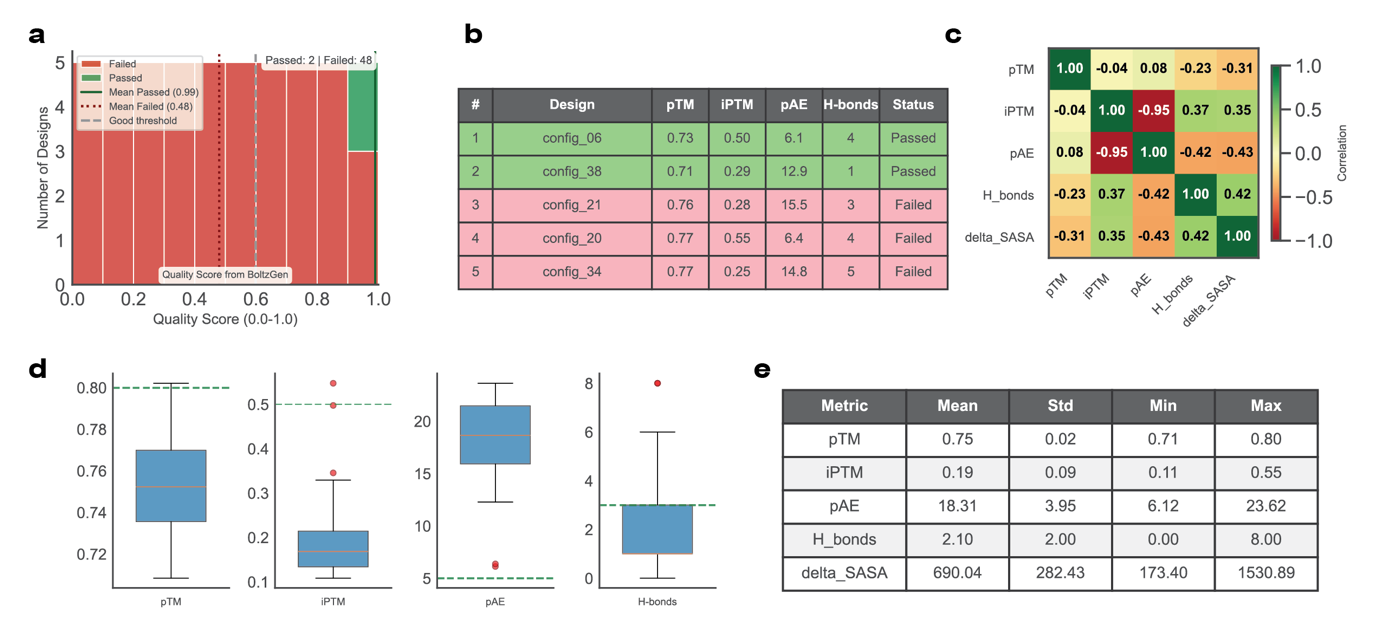

**Figure S4 | Detailed Analysis of Nanobody Design Metrics and Selection Criteria in ProteinMCP.** This figure provides a detailed statistical and comparative analysis of the metrics used to evaluate the de novo designed nanobodies, illustrating the rigorous filtering process. (a) Quality Score Distribution. This histogram plots the distribution of the BoltzGen Quality Score for all 50 generated designs. It demonstrates a clear bimodal distribution, effectively separating the designs into a small group of high-quality candidates that 'Passed' (2 designs, mean score 0.99) and a large majority that 'Failed' (48 designs, mean score 0.48). This highlights the stringent nature of the selection criteria, which is set well above the average quality. (b) Top-Ranked Design Candidates. This table displays the performance metrics for the top five nanobody designs. It quantitatively shows why designs config_06 and config_38 were passed, as they exhibit superior combinations of metrics, such as high interface confidence (ipTM) and low predicted alignment error (pAE), compared to the failed candidates. (c) Inter-Metric Correlation Analysis. This heatmap reveals the relationships between five key performance metrics. A very strong negative correlation of -0.95 between ipTM and pAE is observed, which is a critical indicator that high confidence in the interface structure strongly corresponds to low alignment error. Additionally, moderate positive correlations exist between the number of hydrogen bonds (H_bonds), the change in solvent-accessible surface area (delta_SASA), and the interface confidence (ipTM), suggesting these factors collectively contribute to a favorable binding interface. (d) Statistical Distribution of Design Metrics. This series of box plots visualizes the distribution of the primary metrics across all 50 designs. It shows that while most designs achieve a reasonable overall structure confidence (pTM), the interface confidence (ipTM) and the number of H-bonds are generally low, with only a few successful outliers (red dots) meeting the desired thresholds (green dashed lines). This indicates that achieving a high-quality binding interface is the main challenge in the design process. (e) Summary of Metric Statistics. This table provides a quantitative summary of the distributions shown in panel (d), listing the mean, standard deviation, minimum, and maximum values for each of the five key metrics. This offers a precise statistical foundation for the visual analysis of the design batch's overall performance.

**Table S1 | Summary of MCP tools added in the ProteinMCP.**

| ID | Category | Tool/Method | Primary Function |
| --- | --- | --- | --- |
| 1 | MSA | Online Server (ColabFold) | Rapid MSA generation and structure prediction using the ColabFold server. |
| 2 | MSA | MMseqs2 | High-performance search and clustering for massive sequence sets, enabling deep MSA generation. |
| 3 | Evolution Modeling | PLMC | Infers evolutionary coupling parameters from an MSA for contact and fitness prediction. |
| 4 | Fitness Model | ev+onehot | Predicts protein variant fitness using evolutionary features and one-hot encoding on DMS data. |
| 5 | Fitness Model / Inverse Folding | ESM / ESM-IF | Generates embeddings for fitness prediction and designs sequences for a given backbone structure. |
| 6 | Fitness Model | ProtTrans | Generates protein language model embeddings for predicting fitness, stability, and other functional characteristics. |
| 7 | Solubility Prediction | Protein–Sol | Predicts protein solubility from its amino acid sequence to identify aggregation-prone regions. |
| 8 | Stability Prediction | SpiredStab | Predicts fitness and stability effects of mutations using integrated sequence and structure features. |
| 9 | Mutational Recommendation | MutCompute | Recommends beneficial mutations to enhance protein stability, activity, or binding affinity. |
| 10 | Structure Prediction | AlphaFold3 | Predicts 3D structures of proteins, nucleic acids, and their complexes with atomic accuracy. |
| 11 | Binder Design | BoltzGen | Generates novel protein backbones and sequences for de novo binder design against a target. |
| 12 | Binder Design | BindCraft | Automated de novo design of high-affinity binders using a multi-stage generation and filtering pipeline. |
| 13 | Structure Prediction | Boltz2 | Probabilistic structure prediction that samples alternative conformations and estimates uncertainty. |
| 14 | Inverse Folding | ProteinMPNN | Designs robust and diverse protein sequences for a given backbone structure. |
| 15 | Inverse Folding | LigandMPNN | Designs protein sequences conditioned on a ligand's atomic context to create specific binding pockets. |
| 16 | Structure Relaxing | Rosetta | Refines protein structures through energy minimization to produce physically realistic models. |
| 17 | Structure Prediction | Chai-1 | Predicts protein structure from a single amino acid sequence using a protein language model. |
| 18 | Structure Diffusion | RFDiffusion2 | Generates diverse and novel protein backbones conditioned on specific functional or structural constraints. |
| 19 | Immunogenicity Prediction | NetMHCpan-4.1 | Predicts peptide binding to MHC class I molecules to assess protein immunogenicity. |
| 20 | Immunogenicity Prediction | NetMHCIIpan-4.0 | Predicts peptide binding to MHC class II molecules to assess T-helper cell epitopes. |
| 21 | Structure Prediction | AlphaFold2 | Predicts 3D protein structures from sequence with atomic-level accuracy using deep learning. |
| 22 | Structure Prediction | ESMFold | Provides fast, end-to-end protein structure prediction directly from a single sequence. |
| 23 | MD Simulations | Gromacs | Performs high-performance molecular dynamics simulations to study protein motion and stability. |
| 24 | MD Simulations | Amber | A suite of programs for running molecular dynamics simulations of biomolecules. |

**Table S2 | Summary of public MCP tools supported in the ProteinMCP.**

| ID | Category | Tool/Method | Primary Function |
| --- | --- | --- | --- |
| 1 | Sequence Analysis | Interpro | Provides functional analysis of protein sequences by predicting domains and important sites. |
| 2 | Database | UniProt MCP Server | Programmatic access to the UniProt Knowledgebase for comprehensive protein sequence and functional information. |
| 3 | Database | AlphaFold DB MCP Server | Programmatic access to over 200 million high-quality predicted protein structures. |
| 4 | Database | PDB MCP Server | Enables programmatic search and download of experimentally determined structures from the Protein Data Bank. |
| 5 | Database | STRING DB MCP Server | Access to the STRING database of known and predicted protein-protein interactions. |
| 6 | Database | ProteinAtlas MCP Server | Programmatic access to Human Protein Atlas data on protein expression and localization. |
| 7 | Database | KEGG MCP Server | Automated retrieval of information on metabolic pathways, genes, and genomes from KEGG. |
| 8 | Database | Open Targets MCP Server | Access to the Open Targets platform for validating and prioritizing therapeutic targets. |
| 9 | Database | NCBI Datasets MCP Server | Unified access to retrieve data across all NCBI databases, including genes and proteins. |
| 10 | Literature & Knowledge | Biomedical MCP | Empowers AI agents with specialized biomedical knowledge from authoritative data sources. |
| 11 | Literature & Knowledge | Arxiv MCP | Allows AI agents to programmatically search and access pre-print articles on arXiv. |
| 12 | Literature & Knowledge | BioMCP | Enhances LLMs with protein structure analysis capabilities for more sophisticated structural reasoning. |
| 13 | Literature & Knowledge | PubMed MCP | Enables AI agents to search, access, and analyze over 36 million biomedical articles. |
| 14 | Visualization | PyMOL MCP | Provides a programmatic interface to the PyMOL molecular visualization system for automated image generation. |

**Table S3 | Summary of protein engineering skills supported in the ProteinMCP.**

| ID | Category | Tool/Method | Primary Function |
| --- | --- | --- | --- |
| 1 | Fitness modeling | MSA, MMseqs2, ev+onehot, ESM, ProtTrans | Build and evaluate fitness models on a target dataset |
| 2 | Binder design | BindCraft | Design peptide binders with one of the most significant and widely adopted workflows |
| 3 | Nanobody design | BoltzGen | Design nanobody binders for a target with the highest potential BoltzGen workflow |
